# Sperm production is negatively associated with muscle and sperm telomere length in a highly polyandrous species

**DOI:** 10.1101/2023.03.10.532083

**Authors:** Elisa Morbiato, Silvia Cattelan, Andrea Pilastro, Alessandro Grapputo

## Abstract

Life history theory suggests that aging is one of the costs of reproduction. Accordingly, a higher reproductive allocation is expected to increase the deterioration of both the somatic and the germinal lines through enhanced telomere attrition. In most species, males’ reproductive allocation mainly regards traits that increase mating and fertilization success, i.e. sexually selected traits. In the current study, we tested the hypothesis that a higher investment in sexually selected traits is associated with a reduced telomere length in the guppy (*Poecilia reticulata*), an ectotherm species characterized by strong pre- and postcopulatory sexual selection. We first measured telomere length in both the soma and the sperm over the course of guppy’s lifespan to see if there was any variation in telomere length associated with age. Secondly, we investigated whether a greater expression of pre- and postcopulatory sexually selected traits is linked to shorter telomere length in both the somatic and the sperm germinal lines, and in young and old males. We found that telomeres lengthened with age in the somatic tissue, but there was no age-dependent variation in telomere length in the sperm cells. Telomere length in guppies was significantly and negatively correlated with sperm production in both tissues and life stages considered in this study. Our findings indicate that telomere erosion in male guppies is more strongly associated with their reproductive investment (sperm production) rather than their age, suggesting a trade-off between reproduction and maintenance is occurring at each stage of males’ life in this species.

## Introduction

Telomeres are repetitive sequences of DNA located at the end of the eukaryotic chromosomes (Blackburn, 1991). These noncoding regions assemble with the telomere-binding proteins, and as a single DNA lagging strand complex, form a cap at the end of the chromosome (Armanios & Blackburn, 2012). The telomere complex, on one hand protects the coding sequence from attrition, on the other hand, sets a limitation for the cell replicative potential, therefore acting as a “mitotic clock” (Olovnikov, 1996). Telomeres shorten with each round of somatic cell division because the RNA polymerase is unable to completely replicate the lagging strand, a phenomenon known as the end of replication problem (Watson, 1972). Telomere attrition is not only a byproduct of cell division, but it is also affected by environmental stressors. The telomere sequence is in fact enriched with guanine nucleotides, which are particularly vulnerable to oxidative stress in both somatic and germinal lines (Barnes et al., 2019; Bekaert et al., 2004; Friesen et al., 2020). Regardless of the mechanism involved in telomere erosion, cellular senescence is triggered when telomere length (TL) falls below a certain threshold, the so-called Hayflick limit (Hayflick, 1965). Telomere attrition, on the other hand, can be reduced by the action of the reverse transcriptase telomerase that adds repetitive units of DNA to the telomeric region at each round of cell division (Zvereva et al., 2010). Furthermore, telomere lengthening can rarely occur through recombination between sister telomeres, a mechanism known as alternative lengthening of telomeres (ALT) (Kass-Eisler & Greider, 2000; Liu et al., 2007). Maintaining a balanced TL is critical because abnormal telomere shortening or elongation rates are linked to dysfunctional phenotypes (Vaiserman & Krasnienkov, 2021).

As the rate of telomere attrition affects cellular senescence and death, TL is considered a hallmark of aging (Heidinger et al., 2012). Although there is evidence of a negative correlation between TL and individual longevity in several species, this correlation appears to be weak according to a meta-analysis that compares telomere attrition patterns across non-human vertebrates (Remot et al., 2022). In fact, telomere dynamics (i.e. the rate of TL variation across lifespan) are highly variable among taxa, and wide variability exists also within species and between tissues. Such variation suggests that the rate of telomere erosion does not follow a universal pattern but may rather be shaped by species-specific selective pressure according to life-history strategies (Monaghan & Haussmann, 2006; Olsson et al., 2018; Remot et al., 2022).

The evolution of life-history traits depends on the resource allocation strategy adopted by a population in which a finite budget of resources is expected to be traded-off between growth and reproduction (Kirkwood & Austad, 2000). Evidence that a greater allocation in reproduction is counterbalanced by a shorter longevity has been confirmed by numerous empirical studies which investigated the role of the reproductive load on the aging rate (Candolin, 1998; Hunt et al., 2004; Lemaître et al., 2020; Pike et al., 2007; Preston et al., 2011; Robinson et al., 2006; Van Voorhies & A, 1992). If an increased reproductive investment results in a decreased somatic maintenance and vice versa, individual phenotypic variability should arise consistently with the allocation strategy adopted (Stearns, 1992), including individual variation in TL (Tarka et al., 2018).

The cost of reproduction is expected to be greater in the sex (usually male) that is under stronger sexual selection. Indeed, male competition for the access to the female led to the evolution of costly traits that contribute respectively to increased mating (precopulatory traits, i.e. ornaments and weapons) and fertilization (postcopulatory traits, i.e. ejaculate features) success (Andersson & Simmons, 2006). Allocation in both pre- and postcopulatory traits can come at the cost of higher production of reactive oxygen species (ROS), a source of telomere erosion (Kawanishi & Oikawa, 2004; Monaghan et al., 2009).

As expected, an increased investment in precopulatory traits expression, such as head coloration in the Australian painted dragon, or tail length in the European barn swallow, is negatively correlated with somatic telomere attrition in individuals with more prominent ornamentation (Kauzálová et al., 2022; Rollings et al., 2017). Consistently, TL negatively correlates with male gonad size in the Atlantic silversides (Gao & Munch, 2015) and with sperm velocity in lizards (Friesen et al., 2020), suggesting that the postcopulatory investment leads to a heightened telomere erosion as well. Indeed, enhanced allocation on postcopulatory traits, such as sperm production, can occur through faster spermatogenesis (Ramm et al., 2014). This process implies a higher rate of cellular division, and thus greater telomere erosion in the germinal line, with potential repercussion on fertility (Cariati et al., 2016; Thilagavathi et al., 2013).

Consequently, a trade-off between the investment in sexually selected traits and the rate of telomere erosion is expected to occur in both the somatic and the germinal lines, particularly in species subjected to strong sexual selection, where the investment in sexually selected traits is greater and trade-offs with maintenance (including TL) are more pronounced.

The guppy (*Poecilia reticulata*) is a classical model species for sexual selection studies. Males exhibit orange-colored spots that positively affect male mating success (Houde, 1997). These carotenoid-based ornaments are costly to produce, as indicated by the fact that their expression is condition-dependent (Andersson, 1986; Grether et al., 2004; Locatello et al., 2006; Nicoletto & Kodric-Brown, 1999). Furthermore, the high level of polyandry typical of this species foresees strong sperm competition. Increased sperm production is the most common evolutionary response to increased levels of sperm competition (Lüpold et al., 2020). Guppies make no exception, and the number of sperm transferred during copulation is the best predictor of fertilization success (Boschetto et al., 2011). Male guppies indeed evolved large sperm reserves, that allow them to successfully inseminate several females consecutively (Magris et al., 2020). Once depleted, sperm reserves are restored in a few days (Kuckuck & Greven, 1997). As a result, the ejaculate production likely represents a significant component of the reproductive budget of an individual and shows stronger condition dependence than other sexually selected traits (Devigili et al., 2017; Gasparini et al., 2013). Male guppies show senescence for both male orange colors and ejaculate traits, but ageing is faster for precopulatory traits (Gasparini et al., 2010, 2019).

Here we aim to test whether any variation in TL is associated to either age (telomere dynamics) or pre- and postcopulatory sexually selected traits investment in male guppies. We did the following predictions: i) to observe little variation in the TL of somatic and sperm cells as a function of age in guppies according to previous findings on short-lived ectotherm species (Harel et al., 2015; Lund et al., 2009; Olsson et al., 2018; Remot et al., 2022); and ii) if a higher investment in sexually selected traits is traded-off against maintenance, males with enhanced sexual traits will have shorter telomeres, supporting the trade-off hypothesis (TOH).

We first estimated telomere dynamics in the somatic and in the sperm cells in order to disclose any evidence of somatic or reproductive aging in this species. We thus measured somatic TL at birth (undetermined sex), at 5±1 (full adult males) and at 12±1 months (old males). We further measured sperm TL at 5±1 and 12±1 months. Second, we assessed the relationship between the investment in pre- (relative area of orange spots) and postcopulatory traits (relative sperm production) and TL in the somatic and in the sperm cells. Since a steeper decline in maintenance and reproduction appears with aging (Jones et al., 2014; Nussey et al., 2013), we estimated the relationship between the expression of sexually selected traits and somatic and sperm TL in both young and old males.

## Material and methods

Guppies used in this experiment are descendants of a stock collected from Lower Tacarigua River in Trinidad. They are maintained as a self-sustained population at the Botanical Garden of the University of Padova where the experimental fish have been collected at the fry stage and subsequently acclimatized to the laboratory conditions in stock tanks.

### Experimental design

Fry were haphazardly collected from monitored stock tanks at 1±1 days old to be subsequently euthanized with an overdose of MS222 (Matthews & Varga, 2012) and stored at −20°C in absolute ethanol until DNA extraction. Experimental males (5±1 and 12±1 months old) were selected haphazardly from the stock tanks and stripped to equalize their initial sperm reserves (Gasparini et al., 2009), and then placed into individual tanks (2,5 l). At the age of 5±1 months all males are sexually mature full adult (Magurran, 2005), while at age 12±1 months males show reproductive aging (Gasparini et al., 2019). After 4 days of isolation to allow replenishment of the ejaculate reserves, males have been photographed to assess body size and coloration and then stripped to assess sperm number. After phenotypic traits capture, males were euthanized with an overdose of MS222 and a sample of muscle and sperm were collected (protocol below) and stored in absolute EtOH at −20 °C.

### Precopulatory trait measurement

Male body size and coloration were capture under a ZEISS Stemi 2000-C stereomicroscope through a single sided digital photograph (Canon EOS 450D) after individual anesthesia in MS222 (0.15 g/L) water-based solution (Chambel et al., 2015).

The relative orange area (the orange area on the body area) was measured with ImageJ software (http://rsbweb.nih.gov/ij/download.html) (Evans et al., 2003).

### Postcopulatory trait measurement

Sperm collection and count occurred 4 days after a first strip procedure, a step made with the purpose of flattening sperm reserves variation belonging to putative differences in mating frequencies. After anesthesia, each male was placed on a black slide in 800 μL of saline solution (NaCl 0.9%) under a stereomicroscope. His gonopodium was swung over a 180 degrees angle for 4 times, afterwards, a gentle pressure was applied to the abdomen cavity and sperm were released. Guppies’ sperm are packaged in bundles, each carrying around 22.000 sperm cells (Cattelan et al., 2018), that were photographed and counted with ImageJ software.

The remaining sperm bundles were collected and centrifuged at 5000 rpm in a refrigerated Beckman Coulter microfuge 20R at 4°C for 5 minutes, and the pellet stored in 50 μl of saline solution (NaCl 0.9%) at –20°C until needed.

### RTL measurement

A total of 115 somatic tissues (whole body 1±1 days old =27 and muscle 5±1 =62, 12±1=26 months old), and 74 sperm (age: 5±1=56, 12±1=26) samples were collected. Eleven samples out of the total from the 5±1 age class belong to the same males. In the 12±1 age class, both muscle and sperm tissues belong to the same males.

Genomic DNA was extracted from muscle, and sperm using the Bio Basic EZ-10 Spin Column Genomic Minipreps kit for animal sample according to the manufacturer’s protocol except for sperm samples for which the kit ACL lysis solution was replaced by 300 μl of RTL lysis buffer (Qiagen) and 3 μl of mercaptoethanol. Samples were eluted from the column with either 50 μl (muscle) or 20 μl (sperm) of the kit Elution buffer. DNA quality was checked with a Nanodrop ND-2000 C spectrophotometer (Thermo Scientific, USA) for 260/280 ratio greater than 1.8 and 260/230 ratio greater than 1.9. Quantity was measured with the Qbit (Invitrogen) using the AccuGreen Broad Range dsDNA quantification kit (Biotium). Samples were diluted at the concentration of 2.5 ng/μl. Relative telomere length (RTL) was measured using real time qPCR (Cawthon, 2002), technique that provide telomere quantity as ratio between the telomeric DNA and a single-copy reference gene, here the melanocortin 1 receptor (Monteforte et al., 2020). We used the telomere primers Tel1b and Tel2b (Criscuolo et al., 2009) and the melanocortin 1 receptor MCR1-F and MC1-R primers (Monteforte et al., 2020). Amplification cocktail and protocol were the same as in Monteforte et al. (2020) and run in an Applied Biosystems™ 7500 Real-Time PCR System. Each plate contained three interpolate calibrators and a negative control, all run in triplicates. Baseline and cycle quantification (Cq) values were corrected using the LinRegPCR software ver. 2017.1 (Ruijter et al., 2009). Removal of between-run variation was obtained with Factor qPCR (Ruijter et al., 2015). Relative telomere length was obtained following the equation proposed by Pfaffl (Pfaffl, 2001) as reported in Monteforte et al. (2020). We set the acceptance threshold for amplifications efficiency of 100 ± 20%. Interassay coefficients of variation (CV) were 7.9% for telomere plates and 1.1% for MC1R plates, while intraassay CVs were 1.4% for telomere plates and 0.65% for MC1R plates.

### Statistical analysis

We investigated telomere dynamics throughout guppy’s lifespan in the somatic tissue running a linear regression model (LM) with RTL (log transformed to meet the model assumptions) as the dependent variable and age as the explanatory variable. We tested age classes pairwise comparison running a general contrast between factor levels. To quantify the magnitude of the difference between age classes RTL, we calculated the effect size according to Coehn’s d metric. In order to investigate telomere dynamics in the sperm germinal line we run a LM using sperm RTL (log transformed to meet the model assumptions) as the dependent variable and age as the explanatory variable. To seek any trade-off between sexually selected traits and TL we correlated by mean of generalized linear mixed model (GLMM) the RTL as dependent variable with pre- (relative orange area) and postcopulatory (relative sperm production) traits, age (5±1, 12±1 months) and tissue (muscle and sperm) as fixed predictors. Male identity was entered as random factor to account for repeated measures of TL between tissues within males. Starting from the full model, we removed one by one the least significant predictor to confirm the correlation robustness. The reduced models are presented in the Supporting Information (Table S3). All the analysis were conducted using Matlab version R2018a.

This research comply the ethical requirement and was approved by the Italian health ministry, authorization n. 624/2022-PR.

## Results

### Somatic telomere dynamics

We investigated somatic telomere dynamics over the course of the guppy’s lifespan comparing RTL between newborn, adult, and old males. RTL significantly increased in the somatic line as a function of age (Table 1, Figure 1). Mean RTL at birth was 1.131 (± 0.726 SD). Muscle mean RTL was 1.577 (± 0.908 SD) at 5±1 months and 1.868 (±1.221 SD) at 12±1 months. The pairwise comparison between age classes confirms a significant RTL elongation between 1±1 days old and 12±1 months old (p=0.040), while no significance difference was found between 1±1 and 5±1 ages (p=0.139), and between 5±1 and 12±1 months (p=0.578) (Tables S1, Supporting Information). We measured the effect size between age classes, and we found that RTL effect size between 1±1 and 5±1 (Coehn’s d=-0.242) and between 5±1 and 12±1 (Coehn’s d=-0.288) was small according to Cohen’s indexing d, while the effect size tended to be medium while comparing newborn with old males (1±1 and 12±1, Coehn’s d −0.436) (Tables S1, Supporting Information).

**Table 1.** Results from the linear regression model (LM) in which muscle log RTL (Relative Telomeres Length) was the dependent variable and age (1±1 days old, 5±1 and 12±1 months) was the predictor. Significant terms in bold.

| Muscle RTL |  |  |  |
| --- | --- | --- | --- |
| Fixed factors | Estimate | t | P |
| **Age** | 0.013 | 2.382 | **0.018** |
| **R<sup>2</sup>** |  |  |  |
| 0.047 |  |  |  |

**Figure 1.**
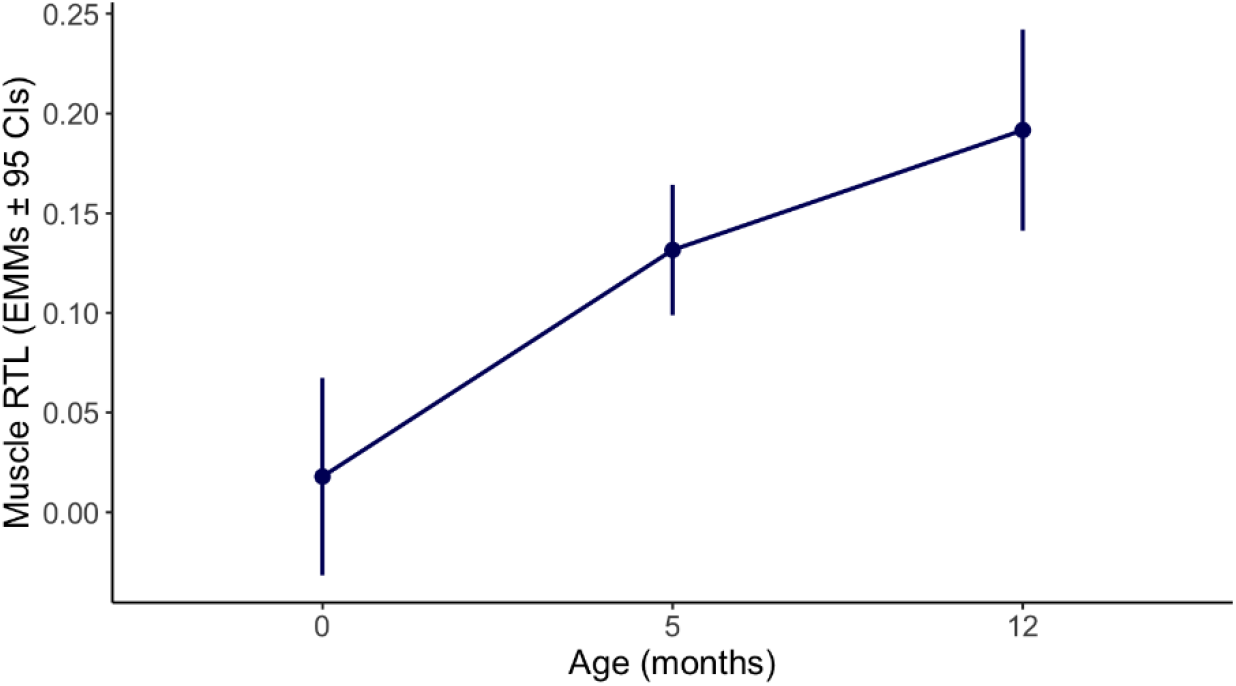
Estimated marginal means of muscle log RTL (Relative Telomeres Length) at different ages (1±1 days old, 5±1 and 12±1 months) from the linear regression model (LM) in Table 1.

### Sperm telomere dynamics

We further analyzed telomere dynamics in sperm of full adult and old males. Sperm mean RTL ranged from 1.549 (±1.1654 SD) at 5±1 months to 1.499 (±1.111 SD) at 12±1 months. No significant variation in sperm RTL was found between adult and old males according to their age (Table 2, Figure 2). The effect size was consistently reporting a very small effect (Cohen’s d=0.044) (Tables S2, Supporting Information).

**Table 2.**
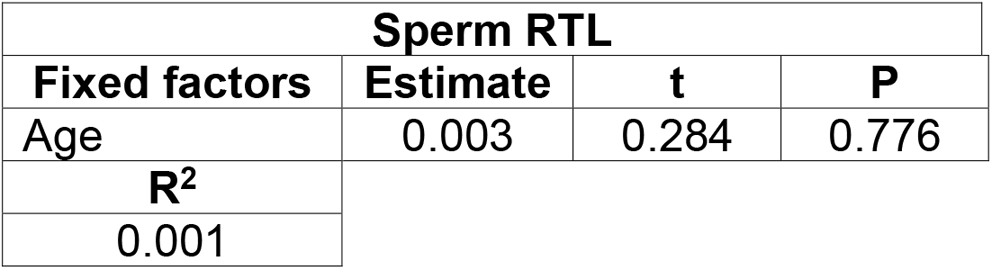
Results from the linear regression model (LM) in which sperm log RTL (Relative Telomeres Length) was the dependent variable and age (5±1 and 12±1 months) was the predictor.

| Sperm RTL |  |  |  |
| --- | --- | --- | --- |
| Fixed factors | Estimate | t | P |
| Age | 0.003 | 0.284 | 0.776 |
| **R<sup>2</sup>** |  |  |  |
| 0.001 |  |  |  |

**Figure 2.**
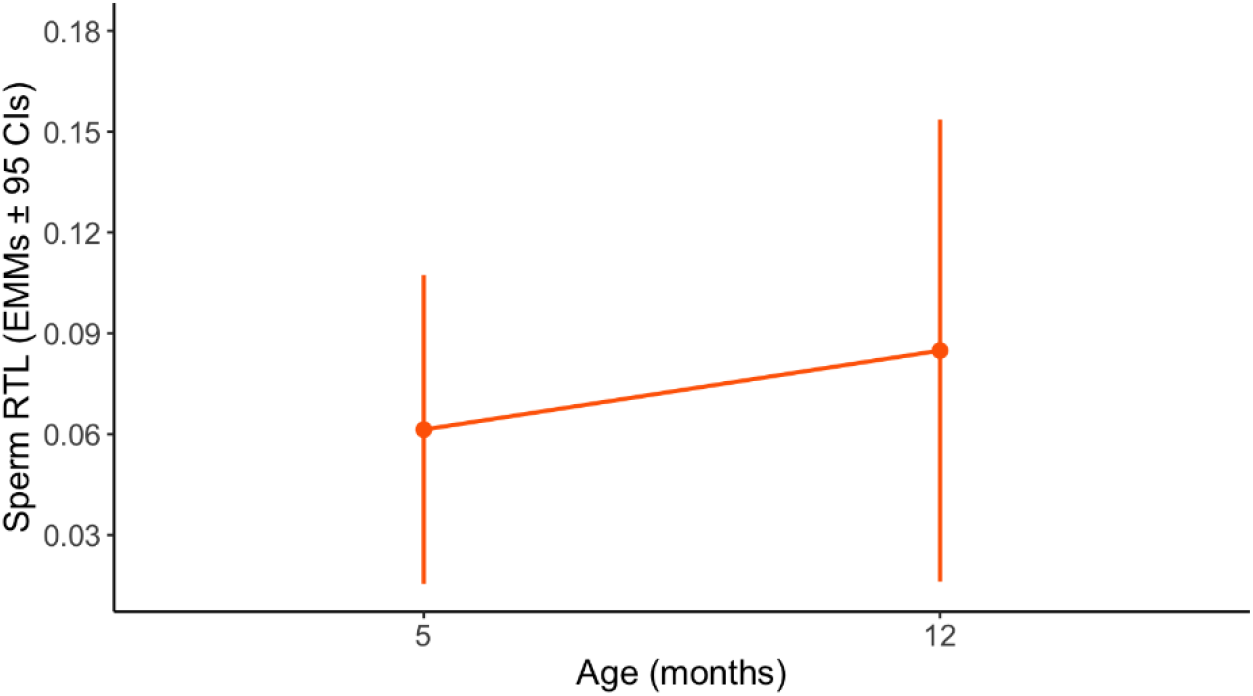
Estimated marginal means of sperm log RTL (Relative Telomeres Length) at different ages (5±1 and 12±1 months) from the linear regression model (LM) in Table 2.

### Sexually selected trait investment and telomere length trade-off

We examined the relationship between somatic and sperm germinal line RTL with the expression of sexually selected traits (relative orange area, relative sperm production) to test whether the investment in pre- and postcopulatory sexually selected traits leads to enhanced telomere erosion. We analyzed this relationship in both adult and old males to consider the putative effect of ageing in telomere attrition. RTL was significantly shorter and negatively correlated with relative sperm production in the somatic and in the sperm germinal lines independently of the age stage (Table 3, Figure 3). The significant association between relative sperm production and RTL persists when removing other predictors from the model (Table S3, Supporting Information). None of the others sexual traits predicted RTL (Table 3). The significant term “tissue” indicates that sperm RTL was consistently shorter than muscle RTL since no significant interaction of the predictors with the tissues was found (Table S4, Supporting Information).

**Table 3.**
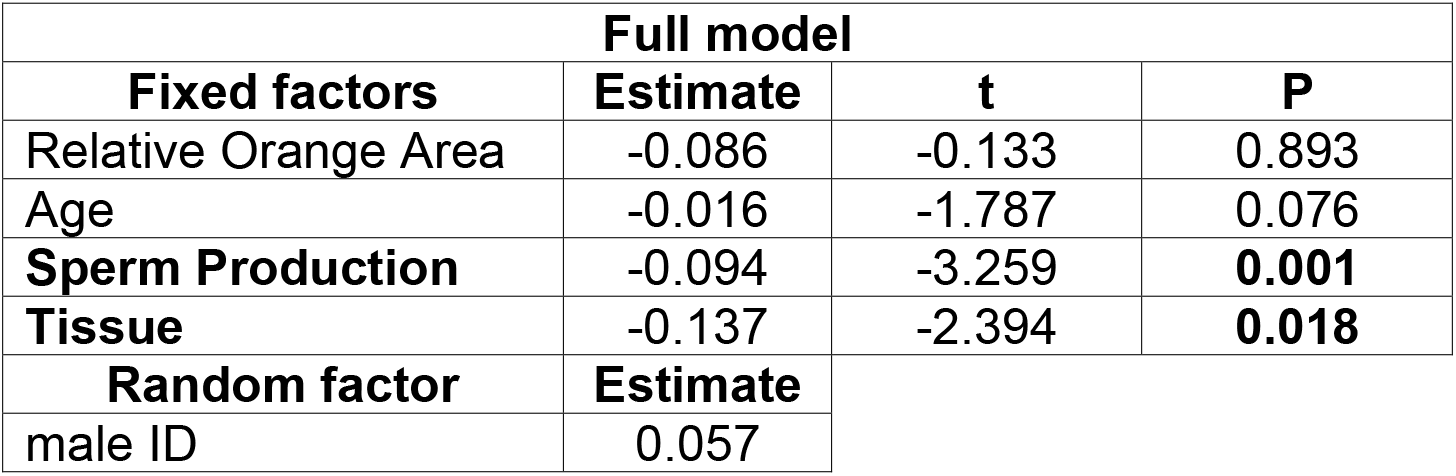
Results from the generalized linear mixed model (GLMM) in which log RTL was the dependent variable and age (5±1 and 12±1 months), relative orange area (orange area on the body area), relative sperm production (standardized residuals of the regression of sperm bundles count on the body area), and tissue (muscle or sperm) were the fixed factors and male ID was the random factor. Significant terms are in bold.

**Figure 3.**
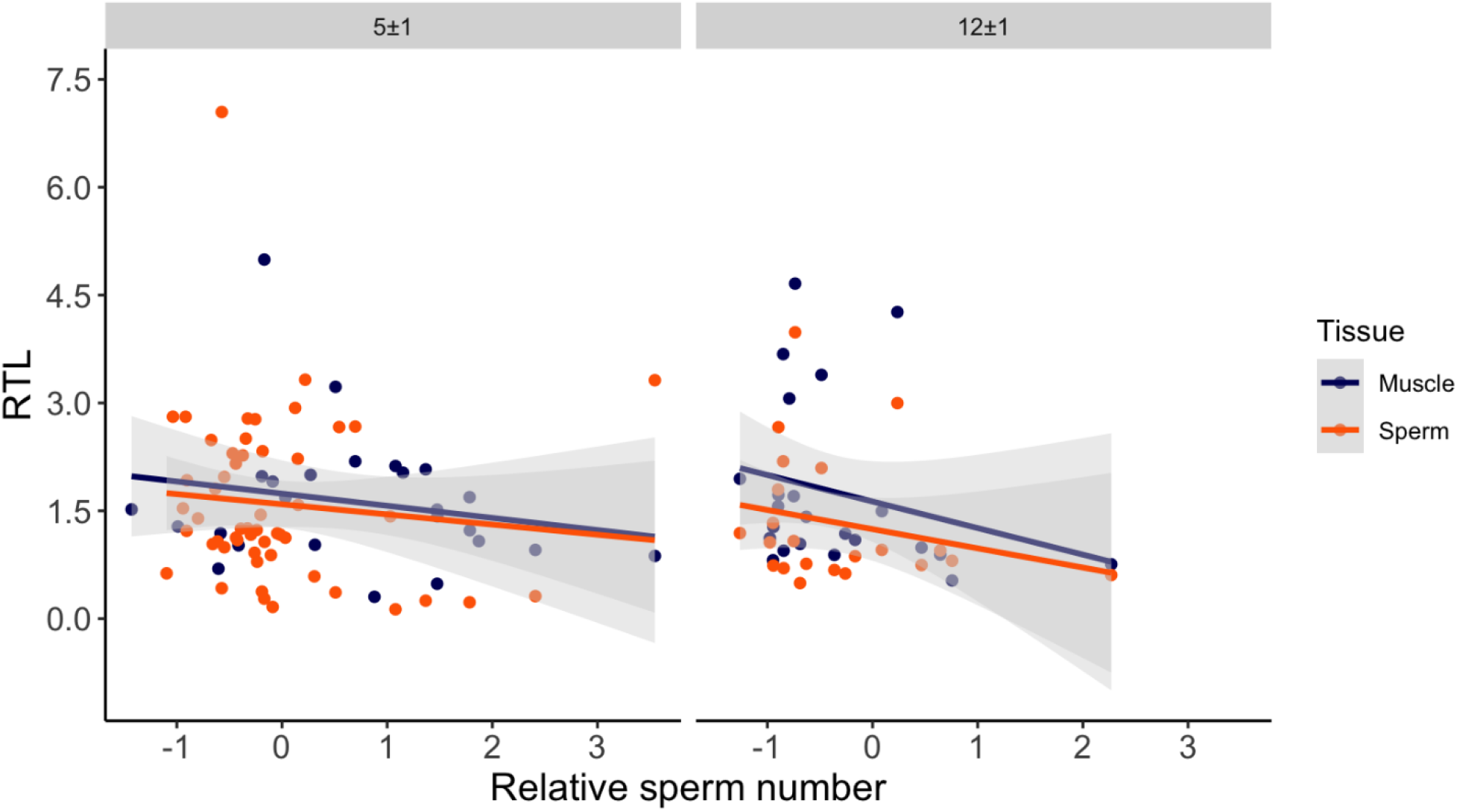
Correlation between log RTL and relative sperm production across ages and tissues. Muscle RTL is reported in blue while sperm RTL is in orange.

## Discussion

### Somatic telomere dynamics

This study aimed to investigate whether the investment in pre- and postcopulatory sexually selected traits (orange coloration and sperm number) is associated with telomere length in both the somatic and the sperm cells. Since the cost of somatic and reproductive maintenance is expected to be greater with aging, we first assessed telomere dynamic in the somatic and sperm cells; secondly, we investigated if any relationship between telomere length and the expression of sexually selected traits was evident in both young and old stage of life in guppy males.

In order to study telomere dynamics, we measured guppy TL at three different ages, namely, at birth (unsexed individuals), 5 (full adult males) and 12 months (senescent males). We found that somatic TL significantly increased with age. There are two possible explanations for this observation. First, in cross-sectional studies (i.e. sampling of different individuals at different ages), increased TL may be the consequence of the selective disappearance of individuals carrying shorter telomeres with aging, hence TL increment as a function of age could be overestimated. Therefore, we cannot exclude a bias due to the selective disappearance of males with shorter telomeres in the old class of age. However, no difference was found in the relationship between TL and age while comparing crosssectional and longitudinal data in a meta-analysis (Remot et al., 2022).

An alternative explanation for the telomere elongation pattern found in guppies relies in the anticancer hypothesis which states that telomerase suppression in senescent stage of life occurs as a mechanism to prevent cancer formation. This suppressed activity of telomerase is proposed to be an evolutionary consequence of higher metabolic and cellular division rates typical of endotherms species with determined growth (Olsson et al., 2018). For this reason, telomeres shortening is less likely observed in small-bodied, short-lived ectotherm species (Olsson et al., 2018; Tian et al., 2018) where the telomerase enzyme is instead consistently expressed throughout lifespan (Anchelin et al., 2013; Lund et al., 2009). However, there is no unique pattern of telomere dynamics among taxa, as telomere elongation over the course of lifespan has been found even in some mammal species (Criscuolo et al., 2018; Hoelzl et al., 2016; Panasiak et al., 2020; Spurgin et al., 2018; Tissier et al., 2022) and no TL variation across ages has been found in fishes (Remot et al., 2022). These results support the idea that telomere dynamics could depend on species-specific life-history strategies rather than following a universal pattern of TL shortening.

### Sperm telomere dynamics

We measured sperm telomere dynamics from the ejaculate of fully adult and old male guppies. We found no statistically significant difference in sperm TL between these two age classes. This result suggests that in guppies, like in other species (Alibardi, 2015; Anchelin et al., 2013; Autexier & Lue, 2006; Harel et al., 2015), the activity of telomerase does not decline with age in proliferative tissues, including male gonads. Indeed, it has been shown that the telomerase is constitutively expressed throughout the lifespan in the gonads of several fish species, such as the lake trout *Salvelinus namaycush* (Purchase et al., 2022), the zebrafish *Danio rerio* (Anchelin et al., 2013), and the killyfish *Nothobranchius furzeri* (Harel et al., 2015).

To date, telomere dynamics in sperm cells have been studied primarily in rodents and humans, where no consistent pattern of positive or negative relationship between TL and age has been found, however, sperm deterioration with age is a common phenomenon (Monaghan & Metcalfe, 2019). Nevertheless, sperm performance (swimming velocity and competitive fertilization success) does not significantly decline with age according to longitudinal studies on guppies (Gasparini et al., 2010, 2019). Coherently, the absence of TL variation in sperm cells across lifespan found in the current study could explain the lack of reproductive aging observed in guppies.

The evolutionary reasons why sperm cell performance age at a slower rate than other male fitness related traits remain unclear (see Gasparini et al. 2019 for a discussion).

### Sexually selected traits investment and telomere length trade-off

In order to reveal a trade-off between TL and male investment in sexually selected traits, we assessed whether males’ orange coloration and sperm production were associated with TL variation in both the somatic and the sperm cells, and in fully adult and senescent males. Our results are strongly indicative of a trade-off between sperm production and telomere attrition in both somatic and sperm cells, and in both fully adult and old males. Intriguingly, the area of orange (carotenoid) coloration did not predict TL in guppy, unlike what has been found in two reptile species (Giraudeau et al., 2016; Rollings et al., 2017).

Collectively, our results indicate that male guppy telomere attrition is more strongly dependent on postcopulatory sexual investment with respect to age. Sperm production, which is strongly condition-dependent in this species (Gasparini et al., 2013), is negatively associated to TL already at 5 months and the same negative association is observed in older males. Considering that sperm production has a high genetic variance and sire heritability (Gasparini et al., 2013), it is possible that this postcopulatory investment is linked to enhanced telomere attrition in both somatic and sperm cells, a hypothesis that could be tested experimentally. The negative relationship between sperm production and TL of both somatic and sperm cells suggests that males with high sperm production may pay two costs: one is a reduced longevity, as shorter somatic TL is expected to be associated with a reduced lifespan. The second cost consists of a reduced sperm competition success, if sperm production is negatively associated with sperm quality (Friesen et al., 2020) and reproductive success (Pauliny et al., 2018).

In conclusion, our findings suggest that male investment in sexually selected traits should be taken into account in the investigation of telomere attrition dynamics since there is increasing evidence that telomere attrition and sexual selection are tightly intertwined (Friesen et al., 2020; Kauzálová et al., 2022; but see Taff & Freeman-Gallant, 2017) and therefore telomere erosion may mediate the cost of reproduction in males. Furthermore, in polyandrous species, male sexual investment and its correlation with TL should be investigated also in postcopulatory traits, especially when they represent a substantial component of male investment in reproduction. Finally, the role of reproduction in telomere dynamics has been so far overlooked, but given its significant contribution to telomeres erosion, it should be included in telomere studies to gain a better understanding of the extended variability in telomere dynamics reported to date.

## Acknowledgements

We thank Gioia Alfonsi e Fabio Palermo for assistance in the molecular data collection. AG was financially supported by the University of Padova (PRID-2020); E.M. and A.P. were supported by the Ministry of University and Research (grant no. 20178T2PSW); AG and AP were supported by the NBFC-PNRR (grant no. C93C22002810006).

## Data availability statement

Additional supporting information may be found online in the Supporting Information section at the end of the article.

## Author contributions

E.M. and A.G. designed the study. E.M. and S.C. collected the samples and the phenotypic data, E.M. and A.G. performed the DNA analysis, E.M; A.P. and A.G. performed the statistical analyses and E.M. wrote the manuscript with contribution from all authors.

## References

Alibardi, L. (2015). Immunolocalization of the telomerase-1 component in cells of the regenerating tail, testis, and intestine of lizards. Journal of Morphology, 276(7), 748–758. https://doi.org/10.1002/jmor.20375

Anchelin, M., Alcaraz-Pérez, F., Martínez, C. M., Bernabé-García, M., Mulero, V., & Cayuela, M. L. (2013). Premature aging in telomerase-deficient zebrafish. Disease Models & Mechanisms, 6(5), 1101–1112. https://doi.org/10.1242/dmm.011635

Andersson, M. (1986). Evolution of condition-dependent sex ornaments and mating preferences: Sexual selection based on viability differences. Evolution, 40(4), 804–816. https://doi.org/10.1111/j.1558-5646.1986.tb00540.x

Andersson, M., & Simmons, L. W. (2006). Sexual selection and mate choice. Trends in Ecology & Evolution, 21(6), 296–302. https://doi.org/10.1016/j.tree.2006.03.015

Armanios, M., & Blackburn, E. H. (2012). The telomere syndromes. Nature Reviews Genetics, 13(10), Article 10. https://doi.org/10.1038/nrg3246

Autexier, C., & Lue, N. F. (2006). The structure and function of telomerase reverse transcriptase. Annual Review of Biochemistry, 75(1), 493–517. https://doi.org/10.1146/annurev.biochem.75.103004.142412

Barnes, R. P., Fouquerel, E., & Opresko, P. L. (2019). The impact of oxidative DNA damage and stress on telomere homeostasis. Mechanisms of Ageing and Development, 177, 37–45. https://doi.org/10.1016/j.mad.2018.03.013

Bekaert, S., Derradji, H., & Baatout, S. (2004). Telomere biology in mammalian germ cells and during development. Developmental Biology, 274(1), 15–30. https://doi.org/10.1016/j.ydbio.2004.06.023

Blackburn, E. H. (1991). Structure and function of telomeres. Nature, 350(6319), Article 6319. https://doi.org/10.1038/350569a0

Boschetto, C., Gasparini, C., & Pilastro, A. (2011). Sperm number and velocity affect sperm competition success in the guppy (*Poecilia reticulata*). Behavioral Ecology and Sociobiology, 65(4), 813–821. https://doi.org/10.1007/s00265-010-1085-y

Candolin, U. (1998). Reproduction under predation risk and the trade–off between current and future reproduction in the threespine stickleback. Proceedings of the Royal Society of London. Series B: Biological Sciences, 265(1402), 1171–1175. https://doi.org/10.1098/rspb.1998.0415

Cariati, F., Jaroudi, S., Alfarawati, S., Raberi, A., Alviggi, C., Pivonello, R., & Wells, D. (2016). Investigation of sperm telomere length as a potential marker of paternal genome integrity and semen quality. Reproductive BioMedicine Online, 33(3), 404–411. https://doi.org/10.1016/j.rbmo.2016.06.006

Cattelan, S., Di Nisio, A., & Pilastro, A. (2018). Stabilizing selection on sperm number revealed by artificial selection and experimental evolution. Evolution, 72(3), 698–706. https://doi.org/10.1111/evo.13425

Cawthon, R. M. (2002). Telomere measurement by quantitative PCR. Nucleic Acids Research, 30(10), e47. https://doi.org/10.1093/nar/30.10.e47

Chambel, J., Pinho, R., Sousa, R., Ferreira, T., Baptista, T., Severiano, V., Mendes, S., & Pedrosa, R. (2015). The efficacy of MS-222 as anaesthetic agent in four freshwater aquarium fish species. Aquaculture Research, 46(7), 1582–1589. https://doi.org/10.1111/are.12308

Criscuolo, F., Bize, P., Nasir, L., Metcalfe, N. B., Foote, C. G., Griffiths, K., Gault, E. A., & Monaghan, P. (2009). Real-time quantitative PCR assay for measurement of avian telomeres. Journal of Avian Biology, 40(3), 342–347. https://doi.org/10.1111/j.1600-048X.2008.04623.x

Criscuolo, F., Smith, S., Zahn, S., Heidinger, B. J., & Haussmann, M. F. (2018). Experimental manipulation of telomere length: Does it reveal a corner-stone role for telomerase in the natural variability of individual fitness? Philosophical Transactions of the Royal Society B: Biological Sciences, 373(1741), 20160440. https://doi.org/10.1098/rstb.2016.0440

Devigili, A., Belluomo, V., Locatello, L., Rasotto, M. B., & Pilastro, A. (2017). Postcopulatory cost of immune system activation in *Poecilia reticulata*. Ethology Ecology & Evolution, 29(3), 266–279. https://doi.org/10.1080/03949370.2016.1152305

Evans, J. P., Zane, L., Francescato, S., & Pilastro, A. (2003). Directional postcopulatory sexual selection revealed by artificial insemination. Nature, 421(6921), Article 6921. https://doi.org/10.1038/nature01367

Friesen, C. R., Rollings, N., Wilson, M., Whittington, C. M., Shine, R., & Olsson, M. (2020). Covariation in superoxide, sperm telomere length and sperm velocity in a polymorphic reptile. Behavioral Ecology and Sociobiology, 74(6), 74. https://doi.org/10.1007/s00265-020-02855-8

Gao, J., & Munch, S. B. (2015). Does reproductive investment decrease telomere length in *Menidia menidia?* PLOS ONE, 10(5), e0125674. https://doi.org/10.1371/journal.pone.0125674

Gasparini, C., Devigili, A., Dosselli, R., & Pilastro, A. (2013). Pattern of inbreeding depression, condition dependence, and additive genetic variance in Trinidadian guppy ejaculate traits. Ecology and Evolution, 3(15), 4940–4953. https://doi.org/10.1002/ece3.870

Gasparini, C., Devigili, A., & Pilastro, A. (2019). Sexual selection and ageing: Interplay between pre-and post-copulatory traits senescence in the guppy. Proceedings of the Royal Society B: Biological Sciences, 286(1897), 20182873. https://doi.org/10.1098/rspb.2018.2873

Gasparini, C., Marino, I. A. M., Boschetto, C., & Pilastro, A. (2010). Effect of male age on sperm traits and sperm competition success in the guppy (*Poecilia reticulata*). Journal of Evolutionary Biology, 23(1), 124–135. https://doi.org/10.1111/j.1420-9101.2009.01889.x

Gasparini, C., Peretti, A. V., & Pilastro, A. (2009). Female presence influences sperm velocity in the guppy. Biology Letters, 5(6), 792–794. https://doi.org/10.1098/rsbl.2009.0413

Giraudeau, M., Friesen, C. R., Sudyka, J., Rollings, N., Whittington, C. M., Wilson, M. R., & Olsson, M. (2016). Ageing and the cost of maintaining coloration in the Australian painted dragon. Biology Letters, 12(7), 20160077. https://doi.org/10.1098/rsbl.2016.0077

Grether, G. F., Kasahara, S., Kolluru, G. R., & Cooper, E. L. (2004). Sex–specific effects of carotenoid intake on the immunological response to allografts in guppies (*Poecilia reticulata*). Proceedings of the Royal Society of London. Series B: Biological Sciences, 271(1534), 45–49. https://doi.org/10.1098/rspb.2003.2526

Harel, I., Benayoun, B. A., Machado, B., Singh, P. P., Hu, C.-K., Pech, M. F., Valenzano, D. R., Zhang, E., Sharp, S. C., Artandi, S. E., & Brunet, A. (2015). A platform for rapid exploration of aging and diseases in a naturally short-lived vertebrate. Cell, 160(5), 1013–1026. https://doi.org/10.1016/j.cell.2015.01.038

Hayflick, L. (1965). The limited in vitro lifetime of human diploid cell strains. Experimental Cell Research, 37(3), 614–636. https://doi.org/10.1016/0014-4827(65)90211-9

Heidinger, B. J., Blount, J. D., Boner, W., Griffiths, K., Metcalfe, N. B., & Monaghan, P. (2012). Telomere length in early life predicts lifespan. Proceedings of the National Academy of Sciences, 109(5), 1743–1748. https://doi.org/10.1073/pnas.1113306109

Hoelzl, F., Smith, S., Cornils, J. S., Aydinonat, D., Bieber, C., & Ruf, T. (2016). Telomeres are elongated in older individuals in a hibernating rodent, the edible dormouse (*Glis glis*). Scientific Reports, 6(1), Article 1. https://doi.org/10.1038/srep36856

Houde, A. E. (1997). Sex, color, and mate choice in guppies. https://press.princeton.edu/titles/6156.html

Hunt, J., Brooks, R., Jennions, M. D., Smith, M. J., Bentsen, C. L., & Bussière, L. F. (2004). High-quality male field crickets invest heavily in sexual display but die young. Nature, 432(7020), Article 7020. https://doi.org/10.1038/nature03084

Jones, O. R., Scheuerlein, A., Salguero-Gómez, R., Camarda, C. G., Schaible, R., Casper, B. B., Dahlgren, J. P., Ehrlén, J., García, M. B., Menges, E. S., Quintana-Ascencio, P. F., Caswell, H., Baudisch, A., & Vaupel, J. W. (2014). Diversity of ageing across the tree of life. Nature, 505(7482), Article 7482. https://doi.org/10.1038/nature12789

Kass-Eisler, A., & Greider, C. W. (2000). Recombination in telomere-length maintenance. Trends in Biochemical Sciences, 25(4), 200–204. https://doi.org/10.1016/S0968-0004(00)01557-7

Kauzálová, T., Tomášek, O., Mulder, E., Verhulst, S., & Albrecht, T. (2022). Telomere length is highly repeatable and shorter in individuals with more elaborate sexual ornamentation in a short-lived passerine. Molecular Ecology, 31(23), 6172–6183. https://doi.org/10.1111/mec.16397

Kawanishi, S., & Oikawa, S. (2004). Mechanism of telomere shortening by oxidative stress. Annals of the New York Academy of Sciences, 1019(1), 278–284. https://doi.org/10.1196/annals.1297.047

Kirkwood, T. B. L., & Austad, S. N. (2000). Why do we age? Nature, 408(6809), Article 6809. https://doi.org/10.1038/35041682

Kuckuck, C., & Greven, H. (1997). Notes on the mechanically stimulated discharge of spermiozeugmata in the guppy, *Poecilia reticulata:* A quantitative approach. Zeitschrift Fur Fischkunde, 4, 73–88.

Lemaître, J.-F., Gaillard, J.-M., & Ramm, S. A. (2020). The hidden ageing costs of sperm competition. Ecology Letters, 23(11), 1573–1588. https://doi.org/10.1111/ele.13593

Liu, L., Bailey, S. M., Okuka, M., Muñoz, P., Li, C., Zhou, L., Wu, C., Czerwiec, E., Sandler, L., Seyfang, A., Blasco, M. A., & Keefe, D. L. (2007). Telomere lengthening early in development. Nature Cell Biology, 9(12), Article 12. https://doi.org/10.1038/ncb1664

Locatello, L., Rasotto, M. B., Evans, J. P., & Pilastro, A. (2006). Colourful male guppies produce faster and more viable sperm. Journal of Evolutionary Biology, 19(5), 1595–1602. https://doi.org/10.1111/j.1420-9101.2006.01117.x

López-Sepulcre, A., Gordon, S. P., Paterson, I. G., Bentzen, P., & Reznick, D. N. (2013). Beyond lifetime reproductive success: The posthumous reproductive dynamics of male Trinidadian guppies. Proceedings of the Royal Society of London B: Biological Sciences, 280(1763), 20131116. https://doi.org/10.1098/rspb.2013.1116

Lund, T. C., Glass, T. J., Tolar, J., & Blazar, B. R. (2009). Expression of telomerase and telomere length are unaffected by either age or limb regeneration in *Danio rerio*. PLOS ONE, 4(11), e7688. https://doi.org/10.1371/journal.pone.0007688

Lüpold, S., de Boer, R. A., Evans, J. P., Tomkins, J. L., & Fitzpatrick, J. L. (2020). How sperm competition shapes the evolution of testes and sperm: A meta-analysis. Philosophical Transactions of the Royal Society B: Biological Sciences, 375(1813), 20200064. https://doi.org/10.1098/rstb.2020.0064

Magris, M., Zanata, I., Rizzi, S., Cattelan, S., & Pilastro, A. (2020). Trade-offs of strategic sperm adjustments and their consequences under phenotype–environment mismatches in guppies. Animal Behaviour, 166, 171–181. https://doi.org/10.1016/j.anbehav.2020.06.016

Magurran. (2005). Evolutionary ecology: The Trinidadian guppy. Oxford University Press.

Matthews, M., & Varga, Z. M. (2012). Anesthesia and euthanasia in zebrafish. ILAR Journal, 53(2), 192–204. https://doi.org/10.1093/ilar.53.2.192

Monaghan, P., & Haussmann, M. F. (2006). Do telomere dynamics link lifestyle and lifespan? Trends in Ecology & Evolution, 21(1), 47–53. https://doi.org/10.1016/j.tree.2005.11.007

Monaghan, P., & Metcalfe, N. B. (2019). The deteriorating soma and the indispensable germline: Gamete senescence and offspring fitness. Proceedings of the Royal Society B: Biological Sciences, 286(1917), 20192187. https://doi.org/10.1098/rspb.2019.2187

Monaghan, P., Metcalfe, N. B., & Torres, R. (2009). Oxidative stress as a mediator of life history trade-offs: Mechanisms, measurements and interpretation. Ecology Letters, 12(1), 75–92. https://doi.org/10.1111/j.1461-0248.2008.01258.x

Monteforte, S., Cattelan, S., Morosinotto, C., Pilastro, A., & Grapputo, A. (2020). Maternal predator-exposure affects offspring size at birth but not telomere length in a live-bearing fish. Ecology and Evolution, 10(4), 2030–2039. https://doi.org/10.1002/ece3.6035

Nicoletto, P. F., & Kodric-Brown, A. (1999). The relationship among swimming performance, courtship behavior, and carotenoid pigmentation of guppies in four rivers of Trinidad. Environmental Biology of Fishes, 55(3), 227–235. https://doi.org/10.1023/A:1007587809618

Nussey, D. H., Froy, H., Lemaitre, J.-F., Gaillard, J.-M., & Austad, S. N. (2013). Senescence in natural populations of animals: Widespread evidence and its implications for bio-gerontology. Ageing Research Reviews, 12(1), 214–225. https://doi.org/10.1016/j.arr.2012.07.004

Olovnikov, A. M. (1996). Telomeres, telomerase, and aging: Origin of the theory. Experimental Gerontology, 31(4), 443–448. https://doi.org/10.1016/0531-5565(96)00005-8

Olsson, M., Wapstra, E., & Friesen, C. (2018). Ectothermic telomeres: It’s time they came in from the cold. Philosophical Transactions of the Royal Society B: Biological Sciences, 373(1741), 20160449. https://doi.org/10.1098/rstb.2016.0449

Panasiak, L., Dobosz, S., & Ocalewicz, K. (2020). Telomere dynamics in the diploid and triploid rainbow trout (*Oncorhynchus mykiss*) assessed by Q-FISH analysis. Genes, 11(7), Article 7. https://doi.org/10.3390/genes11070786

Pauliny, A., Miller, E., Rollings, N., Wapstra, E., Blomqvist, D., Friesen, C. R., & Olsson, M. (2018). Effects of male telomeres on probability of paternity in sand lizards. Biology Letters, 14(8), 20180033. https://doi.org/10.1098/rsbl.2018.0033

Pfaffl, M. W. (2001). A new mathematical model for relative quantification in real-time RT– PCR. Nucleic Acids Research, 29(9), e45.

Pike, T. W., Blount, J. D., Bjerkeng, B., Lindström, J., & Metcalfe, N. B. (2007). Carotenoids, oxidative stress and female mating preference for longer lived males. Proceedings of the Royal Society B: Biological Sciences, 274(1618), 1591–1596. https://doi.org/10.1098/rspb.2007.0317

Preston, B. T., Jalme, M. S., Hingrat, Y., Lacroix, F., & Sorci, G. (2011). Sexually extravagant males age more rapidly. Ecology Letters, 14(10), 1017–1024. https://doi.org/10.1111/j.1461-0248.2011.01668.x

Purchase, C. F., Rooke, A. C., Gaudry, M. J., Treberg, J. R., Mittell, E. A., Morrissey, M. B., & Rennie, M. D. (2022). A synthesis of senescence predictions for indeterminate growth, and support from multiple tests in wild lake trout. Proceedings of the Royal Society B: Biological Sciences, 289(1966), 20212146. https://doi.org/10.1098/rspb.2021.2146

Ramm, S. A., Schärer, L., Ehmcke, J., & Wistuba, J. (2014). Sperm competition and the evolution of spermatogenesis. MHR: Basic Science of Reproductive Medicine, 20(12), 1169–1179. https://doi.org/10.1093/molehr/gau070

Remot, F., Ronget, V., Froy, H., Rey, B., Gaillard, J.-M., Nussey, D. H., & Lemaitre, J.-F. (2022). Decline in telomere length with increasing age across nonhuman vertebrates: A meta-analysis. Molecular Ecology, 31(23), 5917–5932. https://doi.org/10.1111/mec.16145

Robinson, M. R., Pilkington, J. G., Clutton-Brock, T. H., Pemberton, J. M., & Kruuk, L. E. B. (2006). Live fast, die young: Trade-offs between fitness components and sexually antagonistic selection on weaponry in Soay sheep. Evolution, 60(10), 2168–2181. https://doi.org/10.1111/j.0014-3820.2006.tb01854.x

Rollings, N., Friesen, C. R., Sudyka, J., Whittington, C., Giraudeau, M., Wilson, M., & Olsson, M. (2017). Telomere dynamics in a lizard with morph-specific reproductive investment and self-maintenance. Ecology and Evolution, 7(14), 5163–5169. https://doi.org/10.1002/ece3.2712

Ruijter, J. M., Ramakers, C., Hoogaars, W. M. H., Karlen, Y., Bakker, O., Hoff, V. D., B, M. J., & Moorman, A. F. M. (2009). Amplification efficiency: Linking baseline and bias in the analysis of quantitative PCR data. Nucleic Acids Research, 37(6), e45–e45. https://doi.org/10.1093/nar/gkp045

Ruijter, J. M., Ruiz Villalba, A., Hellemans, J., Untergasser, A., & van den Hoff, M. J. B. (2015). Removal of between-run variation in a multi-plate qPCR experiment. Biomolecular Detection and Quantification, 5, 10–14. https://doi.org/10.1016/j.bdq.2015.07.001

Spurgin, L. G., Bebbington, K., Fairfield, E. A., Hammers, M., Komdeur, J., Burke, T., Dugdale, H. L., & Richardson, D. S. (2018). Spatio-temporal variation in lifelong telomere dynamics in a long-term ecological study. Journal of Animal Ecology, 87(1), 187–198. https://doi.org/10.1111/1365-2656.12741

Stearns, S. C. (1992). The Evolution of Life Histories. Oxford University Press.

Taff, C. C., & Freeman-Gallant, C. R. (2017). Sexual signals reflect telomere dynamics in a wild bird. Ecology and Evolution, 00, 1–7. https://doi.org/10.1002/ece3.2948

Tarka, M., Guenther, A., Niemelä, P. T., Nakagawa, S., & Noble, D. W. A. (2018). Sex differences in life history, behavior, and physiology along a slow-fast continuum: A meta-analysis. Behavioral Ecology and Sociobiology, 72(8), 132. https://doi.org/10.1007/s00265-018-2534-2

Thilagavathi, J., Kumar, M., Mishra, S. S., Venkatesh, S., Kumar, R., & Dada, R. (2013). Analysis of sperm telomere length in men with idiopathic infertility. Archives of Gynecology and Obstetrics, 287(4), 803–807. https://doi.org/10.1007/s00404-012-2632-8

Tian, X., Doerig, K., Park, R., Qin, A. C. R., Hwang, C., Neary, A., Gilbert, M., Seluanov, A., & Gorbunova, V. (2018). Evolution of telomere maintenance and tumour suppressor mechanisms across mammals. Philosophical Transactions of the Royal Society B: Biological Sciences, 373(1741), 20160443. https://doi.org/10.1098/rstb.2016.0443

Tissier, M. L., Bergeron, P., Garant, D., Zahn, S., Criscuolo, F., & Réale, D. (2022). Telomere length positively correlates with pace-of-life in a sex-and cohort-specific way and elongates with age in a wild mammal. Molecular Ecology, 31(14), 3812–3826. https://doi.org/10.1111/mec.16533

Vaiserman, A., & Krasnienkov, D. (2021). Telomere length as a marker of biological age: State-of-the-art, open issues, and future perspectives. Frontiers in Genetics, 11, 630186. https://doi.org/10.3389/fgene.2020.630186

Van Voorhies, & A, W. (1992). Production of sperm reduces nematode lifespan. Nature, 360(6403), Article 6403. https://doi.org/10.1038/360456a0

Watson, J. D. (1972). Origin of Concatemeric T7DNA. Nature New Biology, 239(94), Article 94. https://doi.org/10.1038/newbio239197a0

Zvereva, M. I., Shcherbakova, D. M., & Dontsova, O. A. (2010). Telomerase: Structure, functions, and activity regulation. Biochemistry. Biokhimiia, 75(13), 1563–1583. https://doi.org/10.1134/s0006297910130055

