## Supplementary Information for "Sperm production is negatively associated with muscle and sperm telomere length in a highly polyandrous species"

**^1^ Department of Biology, University of Padova, Via Ugo Bassi 58/B, 35131 Padova, Italy**

**^2^ Fritz Lipmann Institute – Leibniz Institute on Aging, Beuntenbergstraße 11, 07745 Jena, Germany**

**^3^ National Biodiversity Future Centre, Piazza Marina 61, 90133, Palermo, Italy**

*** Corresponding author**

****

****

****

****

**EM, 0000-0003-2859-0041; SC, 0000-0002-6303-8810; AP, 0000-0001-9803-6308, AG, 0000-0001-9255-4294**

**Supporting Information**

Table S1. Cohen’s d index of effect size between age classes in the muscle RTL (Relative Telomeres Length). Significant terms in bold.

| **Muscle Cohen’s d** | | |
| --- | --- | --- |
| Factor levels | **RTL 0** | **RTL 5** |
| Mean | 1.328 | 1.577 |
| Variance | 1.563 | 0.825 |
| **d** | **Pooled SD** | **Contrast P** |
| -0.242 | 1.022 | 0.139 |
| Factor levels | **RTL 5** | **RTL 12** |
| Mean | 1.577 | 1.868 |
| Variance | 0.825 | 1.497 |
| **d** | **Pooled SD** | **Contrast P** |
| -0.288 | 1.009 | 0.578 |
| Factors levels | **RTL 0** | **RTL 12** |
| Mean | 1.328 | 1.868 |
| Variance | 1.563 | 1.497 |
| **d** | **Pooled SD** | **Contrast P** |
| -0.436 | 1.231 | **0.040** |

SD, standard deviation.

Table S2. Cohen’s d index of effect size between age classes in the sperm RTL (Relative Telomeres Length).

| **Sperm Cohen’s d** | | | |
| --- | --- | --- | --- |
| Factors | **RTL 5** | | **RTL 12** |
| Mean | 1.549 | | 1.499 |
| Variance | 1.358 | | 1.235 |
| **d** | | **Pooled SD** | |
| 0.044 | | 1.148 | |

SD, standard deviation.

Table S3. Results from the generalized linear mixed model (GLMM) in which RTL (Relative Telomeres Length) was the dependent variable and age (5±1 and 12±1 months), relative orange area (orange area on the body area), relative sperm production (standardized residuals of the regression of sperm bundles on the body area), and tissue (muscle or sperm) were the fixed factors and male ID (identity) was the random factor. Full model and intermediate models after backward elimination of the least non-significant terms are shown. Significant terms are in bold.

| **Full model** | | | | |
| --- | --- | --- | --- | --- |
| **Fixed factors** | **Estimate** | **t** | **P** | **AIC** |
| Relative Orange Area | -0.086 | -0.133 | 0.893 | 56.854 |
| Age | -0.016 | -1.787 | 0.076 |  |
| **Sperm Production** | -0.094 | -3.259 | **0.001** |  |
| **Tissue** | -0.137 | -2.394 | **0.018** |  |
| **Random factor** | **Estimate** |  |  |  |
| male ID | 0.057 |  |  |  |
| **Intermediate model 1** | | | | |
| **Fixed factors** | **Estimate** | **t** | **P** | **AIC** |
| Age | -0.015 | -1.792 | 0.075 | 56.902 |
| **Sperm Production** | -0.093 | -3.251 | **0.001** |  |
| **Tissue** | -0.144 | -2.564 | **0.011** |  |
| **Random factor** | **Estimate** |  |  |  |
| male ID | 0.051 |  |  |  |
| **Intermediate model 2** | | | | |
| **Fixed factors** | **Estimate** | **t** | **P** | **AIC** |
| **Sperm Production** | -0.079 | -2.846 | **0.005** | 58.041 |
| **Tissue** | -0.112 | -2.035 | **0.043** |  |
| **Random factor** | **Estimate** |  |  |  |
| male ID | 0.018 |  |  |  |

Table S4. Results from the generalized linear mixed model (GLMM) in which RTL (Relative Telomeres Length) was the dependent variable and age (5±1 and 12±1 months), relative orange area (orange area on the body area), relative sperm production (standardized residuals of the regression of sperm bundles on the body area), and tissue (muscle or sperm) were the fixed factors and male ID (identity) was the random factor. Each fixed predictors were tested against its interaction with the factor “tissue”.

| **Full model*Tissue** | | | | |
| --- | --- | --- | --- | --- |
| **Fixed factors** | **Estimate** | **t** | **P** | **AIC** |
| Relative Orange Area | 0.123 | 0.118 | 0.906 | 61.993 |
| Age | -0.008 | -0.489 | 0.625 |  |
| Sperm Production | -0.065 | -1.523 | 0.130 |  |
| Tissue | -0.013 | -0.069 | 0.944 |  |
| Relative Orange Area*Tissue | -0.372 | -0.283 | 0.777 |  |
| Age*Tissue | -0.010 | -0.524 | 0.601 |  |
| Sperm Production*Tissue | -0.050 | -0.866 | 0.388 |  |
| **Random factor** | **Estimate** |  |  |  |
| male ID | 0.033 |  |  |  |
